# Investigation of anti-SARS CoV-2 multimeric bicyclic peptide inhibitors in a range of pre-clinical therapeutic settings

**DOI:** 10.1101/2025.10.03.680220

**Authors:** Maximilian A. J. Harman, Katherine U. Gaynor, Guido Papa, Simone Pellegrino, Gustavo Arruda Bezerra, Brian McGuinness, Phillip Jeffrey, Paul Beswick, Steven Stanway, Katerine Van Rietschoten, Liuhong Chen, Ieva Drulyte, Jo Herriott, Edyta Kijak, Eduardo Gallardo-Toledo, Lee Tatham, Parul Sharma, Eleanor Bentley, Jo Sharp, Adam Kirby, Andrew Owen, James P. Stewart, Michael J. Skynner

**Author notes:** Address correspondence to Michael J. Skynner, and James P. Stewart. Maximilian A. J. Harman and Katherine U. Gaynor contributed equally to the writing of this work.

## Abstract

The spread of respiratory viruses, such as Influenza and SARS-CoV-2 has presented significant challenges over the last 30 years with few effective therapeutic options available to this day. Bi-cyclic peptides represent a unique, modular, modality in the antiviral armamentarium against future pandemics. This study provides a deeper evaluation of multivalent bi-cyclic (Bicycle^®^) molecule efficacy in several preclinical SARS-CoV-2 challenge settings. We explore both pre-exposure prophylaxis and post-exposure therapeutic settings via subcutaneous and intranasal routes of administration. We contextualize this further in bespoke scenarios of immune compromisation, and viral transmission. Promisingly, in all studies we observe efficacy, significantly reducing infectious viral burden at each study endpoint. These data further support candidacy of Bicycle molecules as a differentiated antiviral therapeutic class in the context of pandemic preparedness.

**Importance:** The COVID-19 pandemic, triggered a rapid wave of innovation, accelerating the delivery of new vaccine technology and anti-viral treatments. In our first paper, we described the discovery and molecular optimization of Bicycle molecules as a novel drug class for the potential treatment of SARS CoV-2. Here, we have performed deeper characterization of these molecules in established animal models that simulate SARS-CoV-2 transmission, testing more convenient delivery routes, such as intra-nasal. The Bicycle molecules demonstrated positive outcomes in each of these studies and suggest that Bicycle molecules, as convenient and effective anti-viral treatments, could be an important addition to help future preparedness against new viral pandemics.

## Introduction

The last 30 years have seen the emergence of numerous new pathogens, leading to several pandemics. These include SARS-CoV (1), MERS-CoV (2), Zika (3), Avian (4) and Swine (5, 6) flu, HIV (7) and most recently SARS-CoV-2 (8). The most concerning pathogens are those which spread through respiratory infection from human to human, as these have the potential to rapidly spread through global human populations. Controlling the spread of these respiratory viruses, such as Orthomyxoviridae, Paramyxoviridae, Picornaviridae, Coronaviridae, Adenoviridae, and Herpesviridae, has proven challenging over the prior ten-year period (9). A recent analysis of Australian pre-COVID-19 patient samples (n>12000) demonstrated that >50 % of respiratory tract infections were caused by Picornaviridae, Influenza A or Respiratory Syncytial Virus (RSV) (10). Lessons from COVID-19, as well as previous epidemics and pandemics, have taught that the first 100 days are crucial to controlling spread of disease, primarily through good diagnostics and therapeutic or vaccine interventions (11). Although the recent SARS-CoV-2 pandemic was mitigated through the timely development of effective vaccine strategies, it is far from certain that this will be effective for future pathogens (12). In addition, vaccine efficacy has proven to be relatively short lived (13), particularly given SARS-CoV-2 spike antigen’s rapid structural remodeling (14–16). There also remains an underserved population of immunocompromised patients with limited protection against moderate to severe disease (17). Therefore, therapeutic intervention in prophylactic and critical-care treatment settings has been widely sought, yet met only with some, limited, success (18–20). Multiple therapeutic approaches are required which will work in parallel to mitigate future pandemics by controlling transmission and treating severe disease (21, 22).

In our previous work, we described how bicyclic peptides could be used in various multivalent formats to achieve potent SARS-CoV-2 neutralization *in vitro* and *in vivo* (23). By rapidly generating high epitope diversity this technology demonstrated its speed of response to a new and uncharacterized pathogen and its plasticity for timely reformatting to combat the emergence of new viral variants (Figure 1A). We also showed *in vitro* anti-viral activity against multiple SARS-CoV-2 variant pseudotypes, including an early Omicron variant. Here, we use SARS-CoV-2 as a model virus to further evaluate the effectiveness of this platform technology to prevent infection. We assess activity of Bicycle molecules in both immunocompetent and immunocompromised models to explore therapeutic opportunities where vaccination may be problematic or of limited utility (24). We interrogated different infection scenarios, including established SARS-CoV-2 infection and using an innovative airborne transmission model, which is likely to be more representative of person-to-person transmission. We assess utility via a range of convenient dosing routes such as intranasal dosing. These data further demonstrate the potential of this technology to tackle novel pathogens, as illustrated by the work described here on SARS-CoV-2, and suggests that it could be rapidly deployed in the event of future pandemics to tackle new priority pathogens.

**Figure 1.**
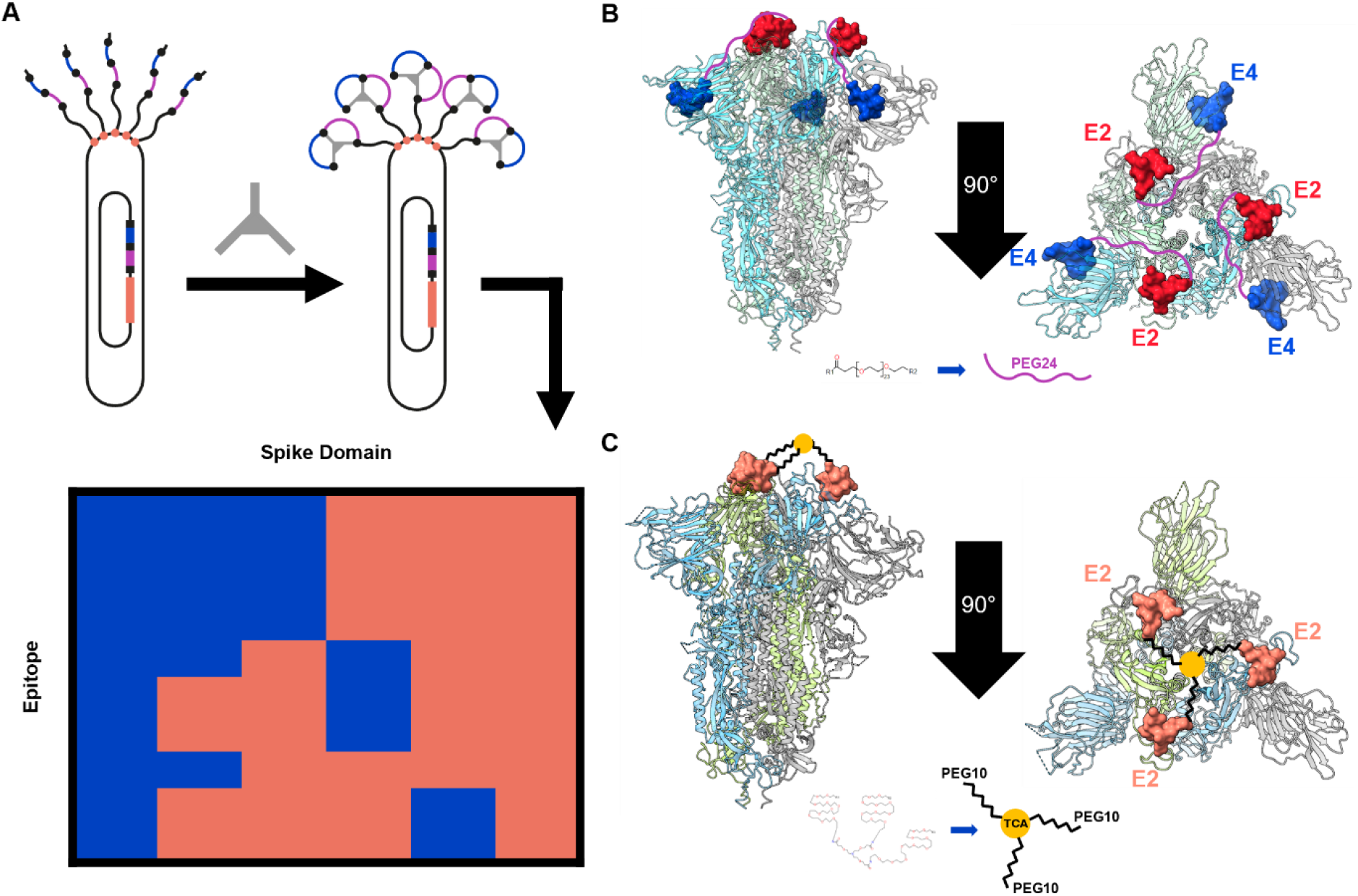
Binders were identified by using bacteriophage libraries to pan the spike protein (adapted from (23)). (A) These libraries displayed linear peptide sequences containing three selectively placed cysteine residues (solid black circles) separating two randomised loops of amino acids (blue and magenta lines). A trivalent chemical scaffold was used to cyclize the linear peptide chains to form the bicyclic peptide moiety. Binders were characterized in a series of biophysical assays to confirm spatially unique epitope clusters. (B) Two Bicycle molecules, E2 and E4, were selected for combination as a biparatopic molecule (named E2E4) separated through C-terminal conjugation to a PEG24 spacer group. Cryo-EM reconstruction (PDB: 9GXG) of the biparatopic E2E4 in complex with the Spike trimer (ribbon) highlights E2 (red surface) bound to the Spike receptor binding domain in surface form, and E4 (blue surface) bound to the Spike N-terminal domain. The PEG24 spacer is shown in magenta as an illustration only. (C) The E2 Bicycle molecule was also combined in a homotrimeric fashion (named E2 Trimer) separated by PEG10 spacers to a trivalent chemical centre. Cryo-EM reconstruction (PDB: 9GXE) of the E2 trimer in complex with the Spike trimer (ribbon) highlights E2 (salmon surface) bound to the Spike receptor binding domain. The PEG10 spacers are depicted by black lines to the yellow trivalent chemical core.

## Results

Two multivalent anti-SARS-CoV-2 Bicycle molecules, previously described as the E2 trimer and the E2E4 biparatopic, exhibited high potency in neutralizing SARS-CoV-2 infection *in vitro* and *in vivo*. The E2 trimer consists of three linked copies of a monomer Bicycle molecule that interact with the SARS-CoV-2 spike (Spike) receptor binding domain (RBD). The monomer subunits are linked via a chemical hinge conjugated to three PEG spacer groups to form a homotrimeric Bicycle ligand. The E2E4 biparatopic consists of the same RBD binding Bicycle molecule (E2), this time conjugated to a PEG spacer group to which is attached a second Bicycle monomer (E4) recognizing the Spike N-terminal domain (NTD). We hypothesized in our previous publication that infection neutralization was due to the ability of these molecules to lock the Spike protein in a pre-fusion format, inhibiting the conformational changes required after engagement of the ACE2 receptor to promote virus infection. To characterize these molecules in complex with Spike protein at molecular level, we used Cryogenic-Electron Microscopy (Cryo-EM) to generate high-resolution reconstructions of Spike protein bound to the E2E4 biparatopic (Figure 1B, Figure S1A and Table S1) and the E2 trimer (Figure 1C, Figure S1B and Table S1). Single particle data processing revealed that for the E2E4 biparatopic, all Spike particles adopted a fully closed conformation (Figure S2). Alternatively, for the E2 trimer, the presence of a partially heterogeneous complex was observed in the sample with 37% of particles adopting a fully closed state and 63% with one “RBD-up” conformation (Figure S3).

The reconstructions obtained for the E2 trimer and the E2E4 biparatopic show the full occupancy of these molecules on the Spike S1 trimeric assembly, respectively on the RBD only and on the RBD and NTD domains. Regarding the E2 trimer, it is important to notice how the scaffold plays a major role in binding, while forming a π-stacking interaction with the side chain of Tyr449. Residues at the end of the first and at the second loop of the Bicycle molecule are the main responsible for binding to the Spike RBD. In more details Norarginine^6^ (Agb^6^), D-Ala^7^ and Thr^8^ (first loop) on the bicycle molecule establish interactions with Leu^452^, Glu^484^, Phe^490^, Gln^493^ and Ser^494^ on Spike. While Leu^9^ and tert-butyl alanine^10^ (tBuAla^10^) interact within a pocket formed by Leu^455^, Phe^456^, Tyr^489^ and Gln^493^ of the target. When we superimpose our structure with the crystal structure of a Bicycle molecule from the same family bound to RBD alone (PDB: 7Z8O) we observe that the binding pose is overall well conserved (RMSD of 0.498 Å of the Cα atomic coordinates). However, the tBuAla^10^ reaches to the backbone of Asn^370^ on the RBD of another protomer of Spike in the current structure, which is generated using the full-length Spike. Structural analysis of the E2E4 biparatopic molecular interactions shows that the binding pose and interactions established of the E2 bicycle on Spike S1 protein is overall highly similar to when crystallized in complex with RBD only (PDB: 7Z8O; RMSD: 0.356 Å at Cα on Bicycle molecule only). The mode of binding of the E4 bicycle to the NTD occurs exclusively through the residues of the two loops, while the scaffold (TATB) is solvent exposed. The Bicycle molecule Ile^1^ and Pro^2^ interact with Phe^175^ and Leu^176^ on Spike, with the latter also involved in stabilizing the peptide’s N-terminally acetylated cysteine. The Bicycle molecule’s Asp^4^ side chain interacts with Spike’s Arg^190^ and His^207^ through the formation of both electrostatic and weak carbon mediated contacts. The peptide’s Trp^5^ fit into a cleft formed between Arg^190^ on one side, and Asn^121^ and Met^177^ on another side of the Spike NTD. Met^7^ on the Bicycle molecule participates in both intra-molecular and inter-molecular interactions, especially with Spike’s Phe^175^. The last two residues of the Bicycle molecule second loop, Ile^8^ and Ala^9^ position within a hydrophobic pocket on Spike, formed by Val^126^, Ile^128^ and Leu^226^. When we superpose the E2 trimer and the E2E4 biparatopic structures we observed good conservation of the binding pose for the E2 Bicycle molecules (RMSD of 0.504 Å of all atoms), suggesting there is no conformational variation promoted by the format of the Bicycle molecule.

Bicycle molecules against Spike protein were efficacious for prophylactic, subcutaneous dosing in hamster SARS-CoV-2 infection models (23). Prophylactic dosing in this way is of inconvenient in pandemic control. Therefore, we sought to explore the therapeutic utility of these compounds, administered subcutaneously, in Golden Syrian Hamster (GSH) models where infection was already established (Figure 2A). Practically, GSHs were infected intranasally with SARS-CoV-2 on day 0 and Bicycle molecules were administered 6h after the infection and then every 8 h for 4 days. All infected groups showed an initial reduction in body weight from day 0 to day 2. However, E2E4-treated GSHs did not show any further reduction compared to the uninfected control. In contrast, body weights of infected untreated GSHs decreased by 7% between day 0 and day 4. On day 4, tissues were removed and analyzed by RT-qPCR (Figure 2C) to assess viral N RNA amplification and plaque assay (Figure 2D) to assess viable SARS-CoV-2 infectivity. E2E4 administration significantly reduced viral RNA in the lung. This reduction was dose-dependent with the most pronounced response at doses of 10 mg/kg. E2E4 treatment also reduced the levels of viable infectious virus particles in the lung homogenate with a statistically significant reduction on day 4 post-infection. In contrast, considerable levels of viable infectious viral particles were detected in the lung samples from untreated GSHs. Collectively, these data indicate that treatment with E2E4 Bicycle molecules can reduce SARS-CoV-2 replication, suggesting a potential therapeutic application following virus exposure.

**Figure 2.**
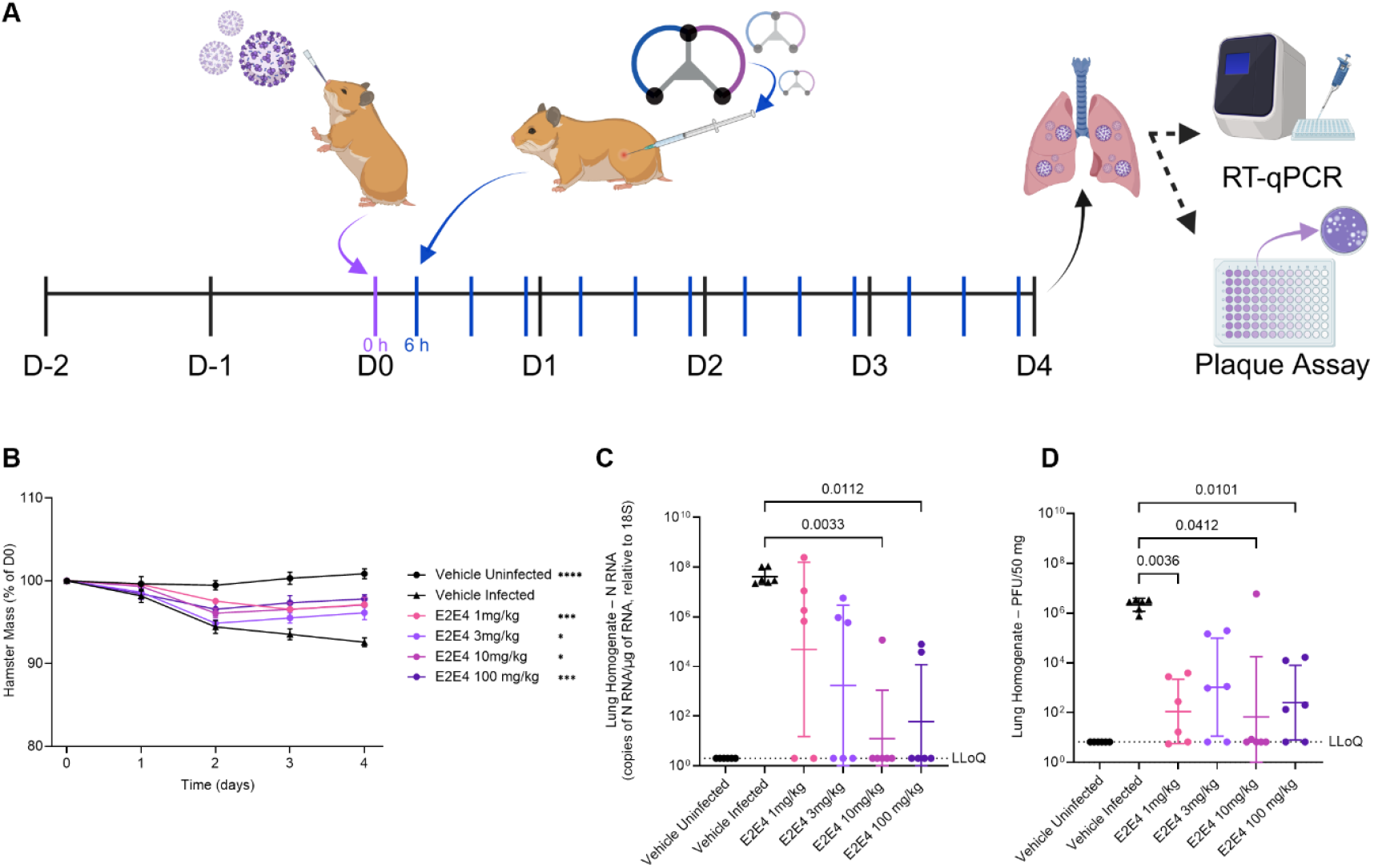
The therapeutic effect of the E2E4 Bicycle molecule was evaluated for in a SARS-CoV-2 challenge model for Golden Syrian Hamsters. (A) Dosing regimen illustrating point of viral challenge (purple), and subsequent subcutaneous administration of E2E4 (blue). At the Day 4 (D4) endpoint, lungs were harvested for homogenization before subsequent RT-qPCR and plaque assays. Created with BioRender.com (B) Hamsters in each treatment group (n = 6) were weighed every day from day 0 to day 4. Body mass is represented as a percentage of the initial body mass on day 0. Two-way ANOVA multiple comparison with Geisser-Greenhouse correction was used to determine statistical significance. * = p ≤ 0.05, ** = p ≤ 0.01, *** = p ≤ 0.001, **** = p ≤ 0.0001. (C) Lung homogenate samples were assayed by RT-qPCR to amplify copies of SARS-CoV-2 nucleocapsid protein mRNA (N RNA). Data presented is normalized as copies of sg N RNA/µg of RNA, relative to 18S ribosomal subunit present in the sample. This is plotted as a geometric mean with 95% confidence intervals. (D) Lung homogenate samples were also tested in plaque assays. This is plotted as a geometric mean with 95% confidence intervals. LLoQ is the lower limit of quantification. Brown-Forsythe and Welch ANOVA with Dunnett’s T3 multiple comparisons test was performed where statistical significance was considered p ≤ 0.05 in all cases.

SARS-CoV-2 is a heightened health risk for immunocompromised patients, who often mount a diminished immune response to vaccination, thereby limiting prophylactic protection (25). To test whether our Bicycle molecules could be beneficial in an immunocompromised setting, we treated GSHs with cyclophosphamide, a potent immunosuppressive agent (100mg/kg) intraperitoneally at day -2, and day 1 (Figure 3A). Only one high dose E2E4 treated group was studied to maintain as high a target coverage at trough concentrations as possible for this proof of principle study using the same dosing regimen as used previously in Figure 2A. Animals showed initial reduction in body weight between day 0 to day 2, which then remained stable until day 4 for uninfected and E2E4-treated GSHs, whilst the body weights of the infected untreated animals continued to decrease by 8% at day 4 (Figure 3B). Here quantification of viral RNA showed a less marked reduction following E2E4-treatment, 670-fold, when compared to immunocompetent animals, and the level of infectious virus was reduced by 16000-fold (Figure 3C and Figure 3D, respectively). These data suggest that, whilst the Bicycle molecules effectively combat the established infection, synergy with the immune system, higher doses or longer treatment duration, may be required to fully clear SARS-CoV-2 infection.

**Figure 3.**
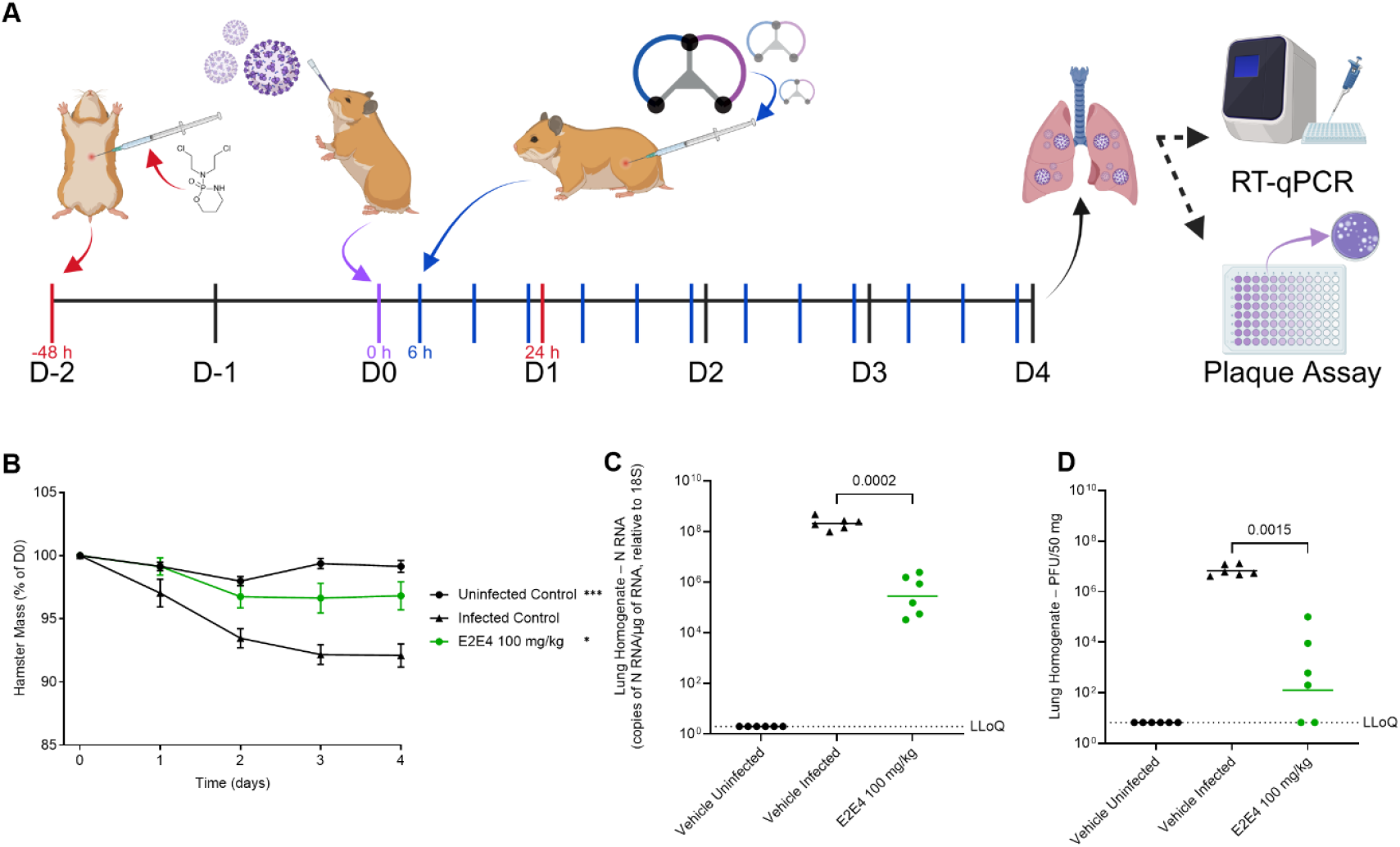
The therapeutic effect for immune suppressed Golden Syrian Hamsters of the E2E4 Bicycle molecule was evaluated in a SARS-CoV-2 challenge model. (A) Dosing regimen illustrating point of viral challenge (purple), cyclophosphamide immune suppression (red), and subsequent subcutaneous administration of E2E4 (blue). At the Day 4 (D4) endpoint, lungs were harvested for homogenization subsequent RT-qPCR and plaque assays. Created with BioRender.com (B) Hamsters in each treatment group (n = 6) were weighed every day from day 0 to day 4. Body mass is represented as a percentage of the initial body mass on day 0. Two-way ANOVA multiple comparison with Geisser-Greenhouse correction was used to determine statistical significance. * = p ≤ 0.05, ** = p ≤ 0.01, *** = p ≤ 0.001, **** = p ≤ 0.0001. (C) Lung homogenate samples were assayed by RT-qPCR to amplify copies of SARS-CoV-2 nucleocapsid protein mRNA (N RNA). Data presented is normalised as copies of sg N RNA/µg of RNA, relative to 18S ribosomal subunit present in the sample. This is plotted as a geometric mean with 95% confidence intervals. (D) Lung homogenate samples were also tested in plaque assays. This is plotted as a geometric mean with 95% confidence intervals. LLoQ is the lower limit of quantification. Brown-Forsythe and Welch ANOVA with Dunnett’s T3 multiple comparisons test was performed where statistical significance was considered p ≤ 0.05 in all cases.

The infection model used in establishing the prophylactic and therapeutic efficacy of E2E4, involved direct administration of virus in solution intranasally. This is not reflective of how SARS-CoV-2 infection occurs in human populations, where airborne virus is inhaled in small droplets. To simulate a more realistic individual to individual model for infection we conducted transmission experiments. We used segregated cages containing a gas permissible divider to separate an infected GSH (Index) from a group of three uninfected GSHs (Contact) (Figure 4A). The presence of a permissible barrier between the infected and uninfected animals allows airborne viral transmission without physical mixing of the two groups. Each GSH “contact” group was studied in duplicate to provide a N=6 sample size per group. The contact vehicle group was exposed to one infected index GSH while the uninfected contact group was exposed to one non-infected index GSH as controls. E2E4 was administered at 100mg/kg subcutaneously in t.i.d. regimen. In the “prophylactic” group, treatment was initiated 2 h pre-exposure to the infected GSH, while, in the “therapeutic” group, treatment was initiated 6 h post-exposure to the index GSH. Quantification of GSH body weight showed no significant change between day 0 and day 4 across all the groups (Figure 4B). On day 4, lung homogenate and nasal turbinate samples of the virus-exposed index GSHs were analyzed by RT-qPCR. High levels of N RNA were detected in both lung and nasal turbinate with the latter showing a higher level, indicating elevated replication in the nasal passage compared to lung (Figure 4C). Oropharyngeal swabs of contact GSHs were also collected on day 1, day 3 and day 4, and N RNA quantified by RT-qPCR (Figure 4D). Between day 1 and day 3, no statistically significant differences in the number of N RNAs copies was seen, however, a drastic reduction was observed at day 4 in prophylactically treated animals. Of note, the infected but untreated group showed an elevated viral burden at day 4 compared to day 1. Therapeutically treated GSHs did not show any statistically significant differences in viral N RNA.

**Figure 4.**
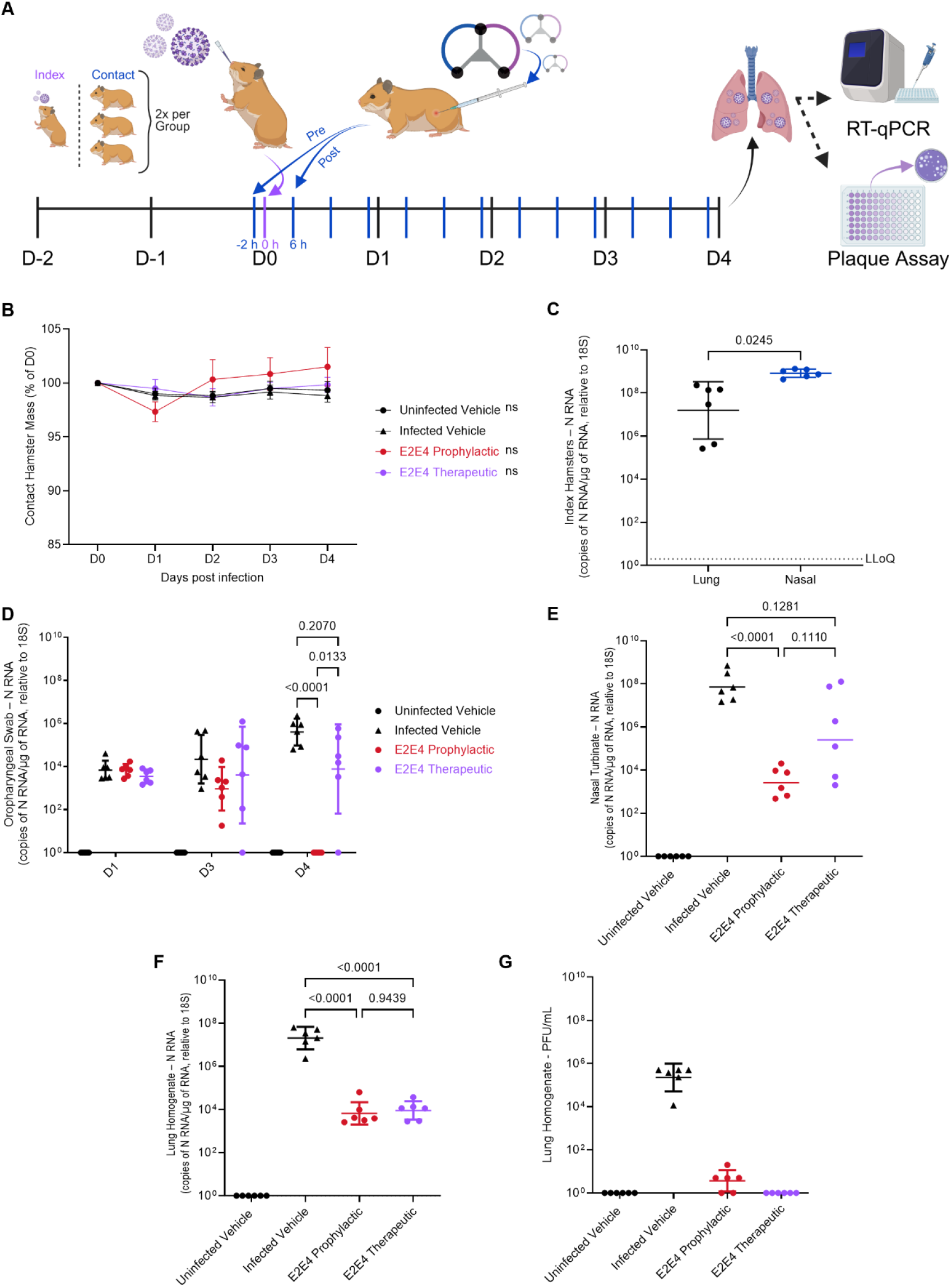
The prophylactic and therapeutic effects of the E2E4 Bicycle molecule were evaluated in a SARS-CoV-2 transmission model for Golden Syrian Hamsters. (A) Dosing regimen illustrating point of viral challenge (purple), and subsequent subcutaneous administration of E2E4 (blue). The grouping of the animals is illustrated on the left-hand side of the figure. Oropharyngeal swabs were taken on day 1 (D1), D3 and D4. At the day 4 (D4) endpoint, lungs were harvested for homogenization as well as nasal turbinate for subsequent assays. Created with BioRender.com (B) Hamsters in each treatment group (n = 6) were weighed every day from day 0 to day 4. Body mass is represented as a percentage of the initial body mass on day 0. Two-way ANOVA multiple comparison with Geisser-Greenhouse correction was used to determine statistical significance. * = p ≤ 0.05, ** = p ≤ 0.01, *** = p ≤ 0.001, **** = p ≤ 0.0001. (C) Index hamster’s lung and nasal samples were assayed by RT-qPCR to amplify copies of SARS-CoV-2 nucleocapsid protein mRNA (N-RNA). Data presented is normalized as copies of sg N RNA/µg of RNA, relative to 18S ribosomal subunit present in the sample. This is plotted as a geometric mean with 95% confidence intervals. Contact hamster oropharyngeal swabs (D), nasal turbinates (E) and lung homogenates (F) were assayed also by RT-qPCR and data is presented as in panel C. (G) Lung homogenate samples were also tested in plaque assays. This is plotted as a geometric mean with 95% confidence intervals. LLoQ is the lower limit of quantification. Brown-Forsythe and Welch ANOVA with Dunnett’s T3 multiple comparisons test was performed where statistical significance was considered p ≤ 0.05 in all cases.

As before, nasal turbinate and lung homogenate samples were collected on termination at day 4 and N RNA quantified by RT-qPCR (Figure 4E and Figure 4F respectively). A highly significant reduction in viral N RNA was observed in nasal turbinate in the prophylactically dosed group when compared to the infected untreated control. A slight decrease in N RNA was detected in the therapeutically dosed group compared to the control, suggesting that administration of Bicycle molecules after viral infection is less effective than pre-treatment using prophylaxis. This is consistent with our previous findings with other SARS-CoV-2 candidate medicines(26, 27).

We also analyzed N RNA amplification in lung homogenates to determine the burden of SARS-CoV-2 in the lower respiratory tract. Both the E2E4 prophylactic and therapeutic groups exhibited a statistically significant reduction compared to the infected vehicle control group. Notably, while the nasal turbinate showed a marked variation between the different animals within a group, the lung homogenates showed less intra-group variations, with both E2E4 treated groups equivalent in their reduction. On day 4 postinfection, lung homogenate was tested for the presence of infectious virus using plaque assays. Treatment by either prophylactic or therapeutic regimen resulted in significant reduction in viral replication when compared to the infected vehicle control group (Figure 3G). Surprisingly, the group treated therapeutically did not show any shedding of infectious viral particles, suggesting that although the viral RNA is not cleared efficiently from the cell, the spreading of the virus across cells is drastically inhibited.

One of the challenges of any antiviral therapeutic is to rapidly deliver the compound such that it reaches the site of action. This was particularly problematic in the early days of the SARS-CoV-2 pandemic when antibody therapies, which lacked oral bioavailability, had to be parenterally administered. As SARS-CoV-2 infection is through respiratory infection, one strategy to prevent infection is through delivery into the nasal/upper respiratory tract. This has the added benefit, that certain therapies can also be absorbed through this route and deliver a level of systemic exposure (Figure S4), which would also be expected to provide additional therapeutic benefit. This route had the added advantage that it is rapid, convenient and needle-free.

To explore the utility of the intra-nasal route of administration, we initially performed pharmacokinetic studies. Here, GSHs were treated with 10 mg/kg of either E2 Trimer or E2E4 biparatopic and sequential blood samples taken over a 24 h period and plasma levels of the compounds determined by LC-MS. (Figure 5A). Both compounds demonstrated similar bioavailability, E2E4 F = 6.1 % and E2 Trimer F = 5.8 % (Figure 5B). Systemic T_1/2_ was also observed between E2E4 at 2.2 h and E2 Trimer at 3.9 h. E2E4 as an empirically smaller molecule approached T_max_ earlier at 0.33 h compared to the larger E2 Trimer at 2.0 h. Given the overall similarity of the PK profiles, 10 mg/kg was determined as an appropriate dose for both compounds using a t.i.d. regimen consistent with our earlier experiments. Due to its larger size and slightly poorer profile the E2 trimer was predicted to maintain IC_90_ coverage at 10 mg/kg for approximately 4 h, however the E2E4 with a slightly better profile was expected to maintain IC_90_ coverage at C_trough_.

**Figure 5.**
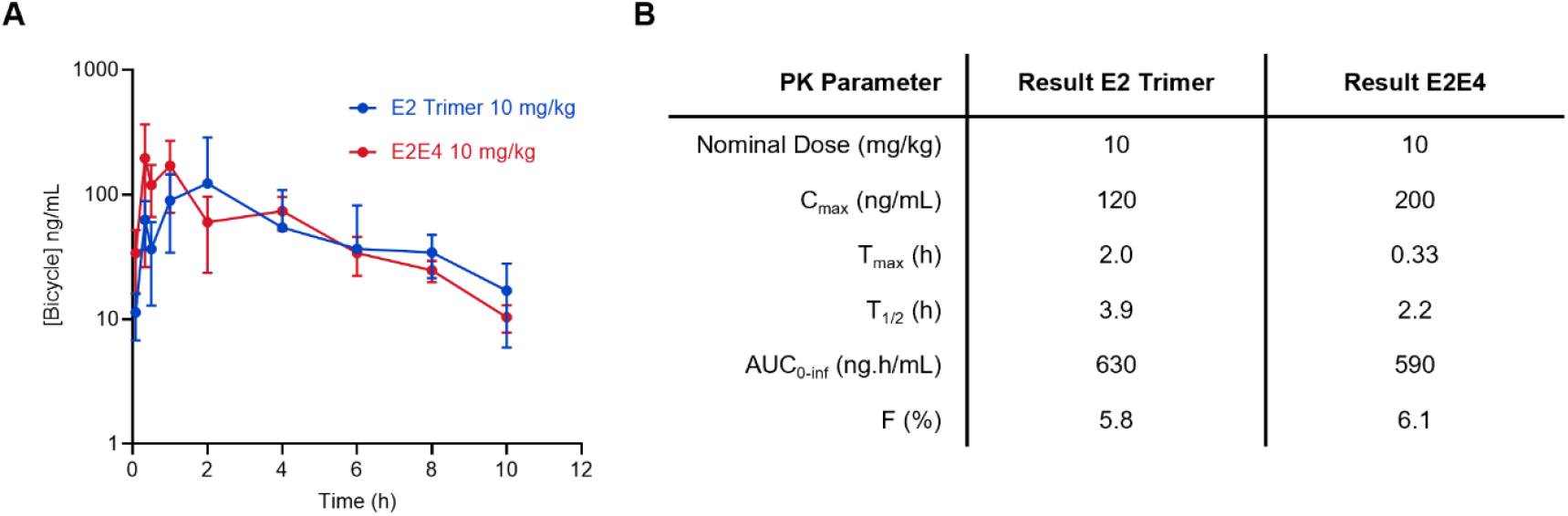
E2E4 and E2 trimer Bicycle molecules were evaluated in an intranasal pharmacokinetic model for Golden Syrian Hamsters. (A) Bicycle multimers were administered (n=6, alternate composite sampling resulting in n=3 per timepoint) via i.n. route of administration. Bicycle plasma concentration over time data displayed as arithmetic mean ± SD for n=3 biological replicates. (B) Pharmacokinetic parameters were subsequently evaluated. Bioavailability (F) was estimated using our previous data studying intravenous administration (23).

To investigate the therapeutic potential of the intra-nasal route of administration, GSHs were treated (10 mg/kg) on a t.i.d. prophylactic regimen, where the animals were challenged with SARS-CoV-2 live virus 1 h after the first dose. (Figure 6A). The infected control group demonstrated a steady decline in GSH body mass over the four-day period losing 9% compared to the Day 0 (pre-challenge) weight. GSH body weight in E2E4 and E2 Trimer groups remained consistent throughout the duration of in-life phase of the experiment (Figure 6B) which was statistically significant when compared to the untreated cohort, E2E4 treated, and E2 Trimer treated groups. Nasal swabs collected every 24 h demonstrated high levels of viral RNA, in both the challenge control and vehicle control groups, 5.3 log_10_(CP/mL) and 5.0 log_10_(CP/mL) respectively (Figure 6C). The E2 Trimer and E2E4 treated groups showed a lower abundance of viral RNA throughout the time course, suggesting that intranasal prophylactic administration of the Bicycle molecules was effective. Nasal turbinate (Figure 6D and 6E) and lung homogenate (Figure 6F and 6G) samples were also collected from all groups at the D4 endpoint to quantify the presence of viral RNA by RT-qPCR and the level of infectious virions using cytopathic plaque assays. Nasal turbinate of both the E2 Trimer and E2E4-treated groups showed significant reductions in viral load when compared to vehicle control. A similar decrease in N RNA copies and infection competent viral particles was observed in lung homogenates in both treatment groups. A slight advantage for the E2E4 over E2 Trimer may be observable in the RT-qPCR data which would reflect the predicted coverage profiles for these molecules at this dose. These results demonstrated initial proof of concept for the delivery of Bicycle molecules as anti-SARS-CoV-2 therapeutics using intranasal administration.

**Figure 6.**
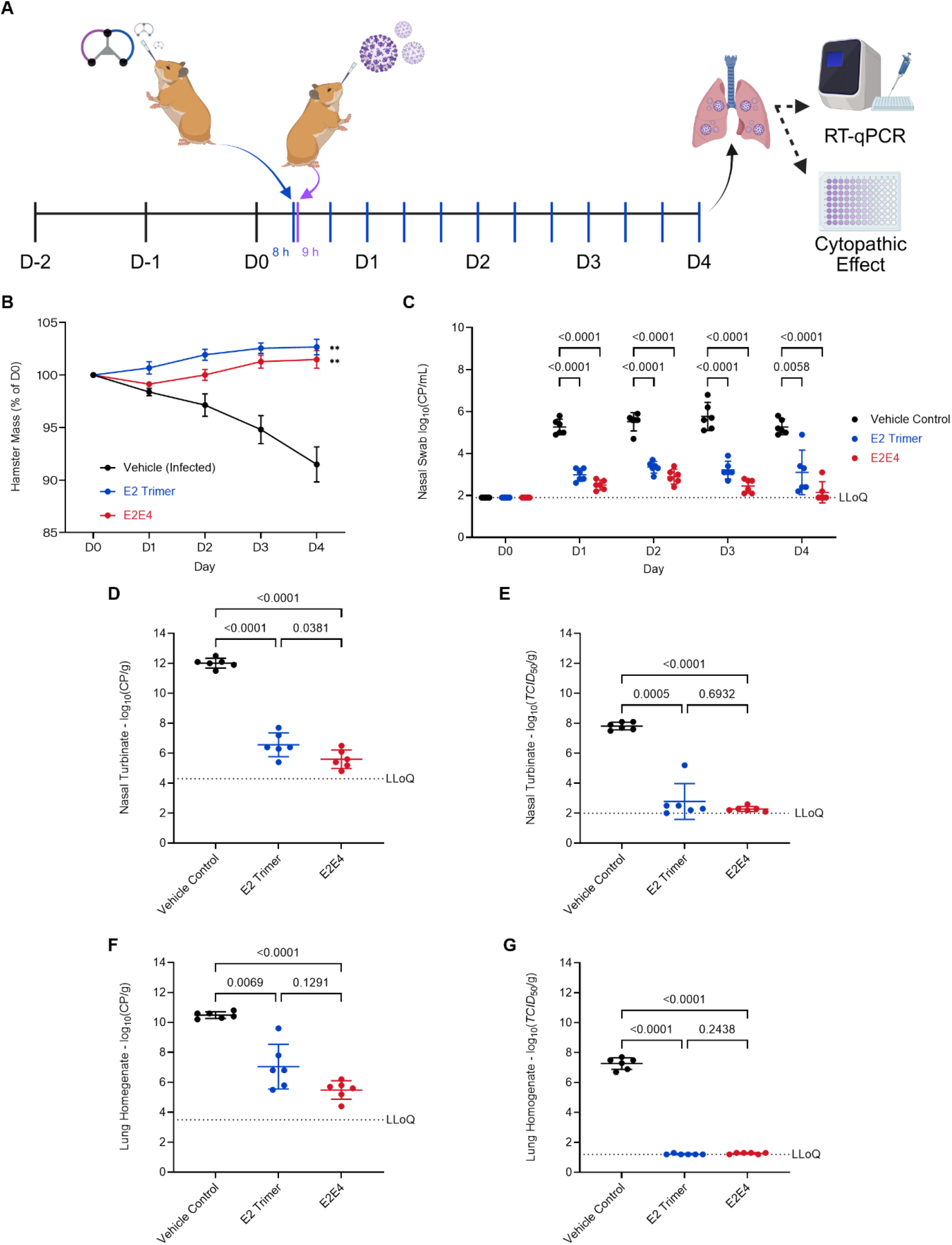
Intranasal prophylaxis of the E2E4 and E2 trimer Bicycle molecules were evaluated in an SARS-CoV-2 challenge model for Golden Syrian Hamsters. (A) Dosing regimen illustrating point of viral challenge (purple), and subsequent subcutaneous administration of E2E4 (blue). Nasal swabs were taken on every day from day 0 (D0) to D4. At the D4 endpoint, lungs were harvested for homogenization as well as nasal turbinate for subsequent assays. Created with BioRender.com (B) Hamsters in each treatment group (n = 6) were weighed every day from D0 to D4. Body mass is represented as a percentage of the initial body mass on D0. Two-way ANOVA multiple comparison with Geisser-Greenhouse correction was used to determine statistical significance. * = p ≤ 0.05, ** = p ≤ 0.01, *** = p ≤ 0.001, **** = p ≤ 0.0001. (C) Nasal swab samples were assayed by RT-qPCR to amplify copies of SARS-CoV-2 envelope protein mRNA (E RNA). Data presented is normalized as copies of E RNA/mL of sample. This is plotted as an arithmetic mean ± SD. Nasal turbinate samples were also assayed by RT-qPCR, as in panel C, however expressed as E RNA/g of sample (D) and for cytopathic effect (E). Cytopathic effect was measured as the TCID_50_/g of sample titrated across a Vero E6 cell monolayer, and data shown as arithmetic mean ± SD. Lung homogenate samples were also assayed in the same way by RT-qPCR (F) and for cytopathic effect (G). LLoQ is the lower limit of quantification. Brown-Forsythe and Welch ANOVA with Dunnett’s T3 multiple comparisons test was performed where statistical significance was considered p ≤ 0.05 in all cases.

## Discussion

We previously described the rapid identification of a “toolkit” of Bicycle molecules that bound and inhibited SARS-CoV-2 infection and how these could be re-configured to respond to mutational change in the virus and to inhibit new viral variants. Primarily, we focused on the chemistry and molecular aspects of inhibition, using prophylactic dosing in animal models. Here, we have looked to now apply these molecules in a more therapeutically relevant setting. Specifically, we demonstrate that these compounds can be administered following viral infection, up to 6 h post infection, in a therapeutic setting, in addition to as prophylactics.

We show that these compounds are effective at neutralising infection via airborne transmission. We also describe that these compounds are effective when administered intranasally, which is potentially a viable strategy that could be used, in conjunction with vaccination, to control the societal speed and spread of disease, as well as a companion therapeutic to treat patients exposed to viral infections. This route likely protects locally at the initial site of infection in the nasal and upper airways, but also through delivery into the systemic circulation likely protection of subsequent viral infection and replication in other tissues and organs. Importantly, we have also looked at one of the patient groups which is still underserved by vaccination, the immune compromised, and show that the same technology can be used to treat viral infections in this vulnerable patient category. Collectively, these data build further confidence that the Bicycle technology platform could be used to tackle future viral threats and form part of the armamentarium for tackling future pandemic threats.

## Materials and Methods

### Cryo-EM grids preparation, data collection and image processing

For preparation of both complexes, E2E4 biparatopic and E2 trimer with Spike S1, 5.4 μl of a 33 μM (initial target stock concentration) SARS-CoV-2 hexaproline S ectodomain was mixed with 3-fold molar excess of bicycle (0.6 μL of a 1mM initial stock concentration) and then incubated for ∼10 min at RT. Immediately before blotting and plunge freezing, fluorinated octyl maltoside (FOM) was added to the sample, resulting in a final FOM concentration of 0.01% (w/v). For both conditions, 3 μL of S-BCY complexes was applied to glow-discharged (20 mA, 30 s; Quorum GloQube) Quantifoil R1.2/1.3 grids (Quantifoil Micro Tools GmbH), blotted for 5 s using blot force 0 and plunge-frozen into liquid ethane using Vitrobot Mark IV (Thermo Fisher Scientific). The data were collected on the Thermo Fisher Scientific Krios G4 Cryo Transmission Electron Microscope equipped with Selectris X Imaging Filter (Thermo Fisher Scientific) and Falcon 4 Direct Electron Detector (Thermo Fisher Scientific) operated in Electron-Event representation mode. Data processing was performed in cryoSPARC (28) single-particle analysis suite. Raw data were imported in cryoSPARC. For the E2E4 biparatopic complex, after Patch motion correction and Patch CTF estimation, 523,554 particles were picked from 10,297 movies. After two rounds of 2D classification, a small subset of particles was used to generate an initial volume by *Ab initio* reconstruction. After this step, 429,402 particles were used to run a homogeneous refinement (28) with C3 symmetry imposed, yielding a cryo-EM map with an overall resolution of 1.92 Å. For the E2 trimer complex, after Patch motion correction and Patch CTF estimation, 472,243 particles were picked from 6,697 movies. After two rounds of 2D classification and an Ab initio reconstruction job, two main classes were identified: fully closed and “one RBD-up” states. For the former 117,262 particles were used to run a homogeneous refinement (28) with C3 symmetry imposed, yielding a cryo-EM map with an overall resolution of 2.3 Å. For the “one RBD-up” volume, 196,044 particles were used to run a homogeneous refinement (28) with C1 symmetry imposed, yielding a cryo-EM map with an overall resolution of 2.4 Å. For both complexes, the “Gold Standard” Fourier shell correlation (FSC) criterion (FSC = 0.143) was used for resolution estimates. Data collection and reconstruction parameters can be found in Table S1. Figures of the reconstructions were generated using ChimeraX (29).

### Model building and validation

UCSF ChimeraX (29) (version 1.9) and Coot (30) (version 0.9.8) were used for model building. As a starting point for modelling the BCY-bound Spike complex, the cryo-EM structure of the SARS-CoV-2-S (PDB ID 6XR8) was initially rigid-body fitted into the density map using the UCSF Chimera “Fit in map” tool (29). The docked model was then mutated to introduce the proline mutation at six different sites, to match the 6P S construct used in this study. The Bicycle molecules were built *de novo* using Coot (30). Iterative rounds of manual fitting and building in Coot and real space refinement in Phenix (31) were carried out to improve the initial model. The final model was validated within Phenix (31), Figures of the models were generated using ChimeraX (29)

### Golden Syrian Hamster Therapeutic Subcutaneous Challenge

Golden Syrian Hamster challenge studies were conducted in accordance with UK Home Office Scientific Procedures Act (ASPA, 1986) and project license PP4715625. Male Golden Syrian hamsters were purchased from (Janvier, 85-110 g). Hamsters were housed in groups of six animals. Photoperiod was 12 h on 12 h off light/dark cycle. Ambient animal room temperature is 21 °C, controlled within ± 2 °C and room humidity 50% ± 5%. On Day 0, hamsters were infected intranasally with 10^4^ PFU/hamster SARS-CoV-2/human/Liverpool/REMRQ0001/2020 using 100 µL inoculum, 50 µL per nostril. Viral inoculum was administered pre the concurrent drug or vehicle (25 mM Histidine HCl buffer, 10% (w/v) sucrose at pH 7.0) dose. At 6 h post infection hamsters were treated by subcutaneous injections three times per 24 h period, every 8 h (t.i.d.). Hamsters continued to be treated t.i.d. with either vehicle or Bicycle molecule until Day 4 post infection. An uninfected control group of hamsters was also added, receiving vehicle only (t.i.d.). E2E4 Bicycle was administered t.i.d. at 100 mg/kg, 10 mg/kg, 3 mg/kg and 1 mg/kg. On Day 4 the animals were sacrificed using an overdose of pentobarbitone. Lung homogenate and nasal turbinate samples were removed for RT-qPCR and plaque assay analysis as described in our previous work (32). Additionally, the same challenge protocol was replicated for immunosuppressed groups; untreated vehicle t.i.d., infected vehicle t.i.d. and 100 mg/kg t.i.d. E2E4. These groups were immunosuppressed using an intraperitoneal injection of 100 mg/kg cyclophosphamide (Baxter Healthcare Ltd) at both Day -2 to pre-infection and Day 1 post-infection. For the Day 1 administration, the treatment condition was administered immediately prior to the immunosuppression condition.

### Golden Syrian Hamster Prophylactic and Therapeutic Subcutaneous Transmission Challenge

Golden Syrian Hamster challenge studies were conducted in accordance with UK Home Office Scientific Procedures Act (ASPA, 1986) and project license PP4715625. Male Golden Syrian hamsters were purchased from (Janvier, 85-110 g). Hamsters were housed in groups of four animals. Photoperiod was 12 h on 12 h off light/dark cycle. Ambient animal room temperature is 21 °C, controlled within ± 2 °C and room humidity 50% ± 5%. On Day 0, six “index” hamsters were infected intranasally with 10^4^ PFU/hamster SARS-CoV-2/human/Liverpool/REMRQ0001/2020 using 100 µL inoculum, 50 µL per nostril. All cohorts had a sample size N=6. The vehicle (25 mM Histidine HCl buffer, 10% (w/v) sucrose at pH 7.0) treated uninfected control cohort was not exposed to infected index hamsters. Infected and uninfected control groups received vehicle 2 h pre contact, and subsequently t.i.d. until Day 4. E2E4 drug was administered t.i.d., until Day 4, to two cohorts, one 2 h pre contact (prophylactic), one 6 h post contact (therapeutic) at 100 mg/kg. Each cohort was split into two groups of three hamsters and housed in a Techniplast GR1800DIV cage on the day of infection (Day 0). This cage has a plastic perforated divider that allowed airflow from one side to the other. Contact was initiated by co-housing one infected hamster in each cage of three treated hamsters, separated by the perforated divider. At Day 4 the animals were sacrificed using an overdose of pentobarbitone. Lung homogenate and nasal turbinate samples were removed for RT-qPCR and plaque assay analysis as described in our previous work (32).

### Golden Syrian Hamster Prophylactic Intranasal Challenge (Viroclinics)

Hamster challenge studies were carried out in the central animal facilities of Viroclinics Biosciences B.V. in Schaijk, The Netherlands, under conditions that meet the standard of Dutch law for animal experimentation and are in agreement with the “Guide for the care and use of laboratory animals”, Institute for Laboratory Animal Research (ILAR) recommendations, AAALAC standards. Ethics approval was made by the local animal welfare body for the present study and registered under number: 27700202114492-WP37. For all animal procedures, the animals were sedated with isoflurane (3-4%/O2). These procedures included subcutaneous and intranasal dosing, blood sampling, challenge, throat swabs and euthanasia.

For this study, male Syrian golden hamsters (Janvier, 89 – 114g) were split into four treatment regimes, containing 6 animals per group: E2 trimer at 10 mg/kg t.i.d.; E2E4 biparatopic at 10mg/kg t.i.d.; Vehicle (25 mM Histidine HCl buffer, 10% (w/v) sucrose at pH 7.0). The treatment protocol was administered via an intranasal route with 50 μL/100 g from Day 0, up to and including Day 3. 1 h post initial treatment, the hamsters were infected intranasally with 10^2^ TCID_50_ BetaCoV/Munich/BavPat1/2020 using 100 µL inoculum, 50 µL per nostril. On Day 4 post infection, animals were euthanized and lungs taken for analysis. During the study, body mass was tracked, and blood samples and throat swabs taken at selected time points. For virological analysis, lung and nose tissue samples were weighed, homogenized in 1.5 mL inoculation medium (DMEM containing L-Glutamine, Penicillin, Streptomycin, Amphotericin B and Fetal Bovine serum, and centrifuged briefly before titration. RT-qPCR and viral cytopathic effect (CPE) analysis was conducted as per our previous work (23).

### Statistical Analysis

Statistical analyses were conducted using GraphPad Prism v10. Body mass data were analyzed using a repeated measures two-way analysis of variance (ANOVA) with Tukey’s multiple comparisons and Geisser-Greenhouse correction, α=0.05. Nasal swab datasets were analysed also by this method. For both RT-qPCR and plaque assay endpoint datasets unpaired Welch’s two-tailed t-tests were used to compare infected control groups with only one additional treatment group. For the same assays, when comparing infected control groups against more than 1 treatment group, a Brown-Forsythe and Welch ANOVA with Dunnett’s T3 multiple comparisons test was performed with α=0.05.

## Acknowledgements

This work was supported by funding from Innovate UK under the grant ID 73786. We would like to thank Viroclinics Xplore for in vivo studies exploring intranasal prophylaxis, WuXi AppTec for intranasal pharmacokinetic studies, and Leo C. James for helpful conversations regarding the planned studies. The authors declare M.A.J.H., K.U.G., G.P., S.P., G. A. B., B.M., P.J., P.B., S.S., K.V.R., L.C. and M.J.S. are employees, and/or shareholders in Bicycle Therapeutics Plc, the parent company of BicycleTx Ltd. These authors may also be listed as inventors on patents or patent applications pertaining to the molecules described.

---

**Figure S1.**
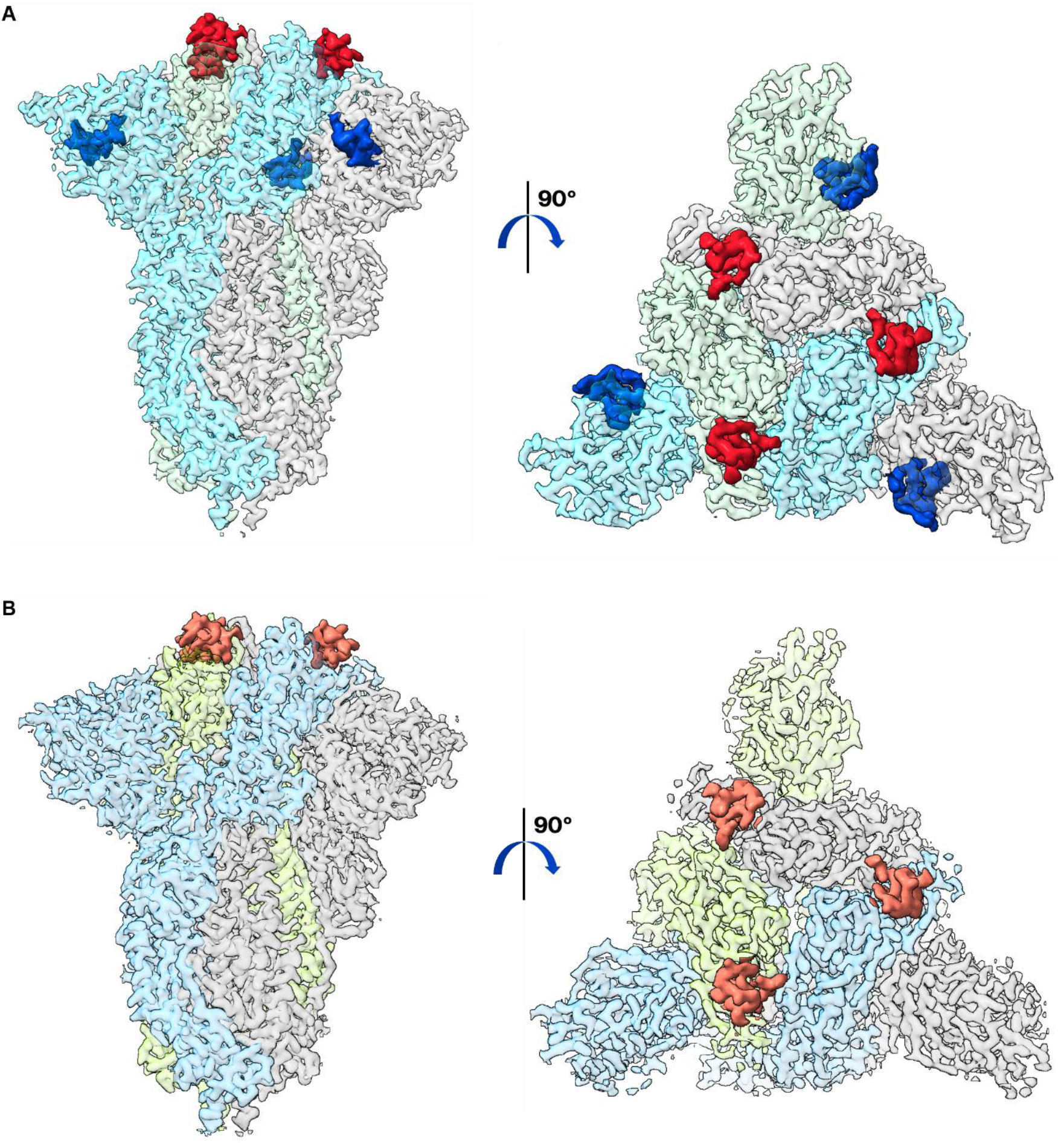
High-resolution cryo-EM reconstructions allow atomic modelling of the E2 trimer and the E2E4 bicycle molecules in complex with SARS-CoV-2 spike antigen. A) Density map representation of the E2E4 biparatopic complex, with each monomer of the spike trimeric assembly coloured differently (cyan, light green and light grey) and the bicycle molecules coloured in red (E2) and blue (E4). Two views, 90 degrees apart, are shown for clarity. B) Density map representation of the E2 trimer complex, with each monomer of the spike trimeric assembly coloured differently (light blue, lime and light grey) and the bicycle molecule coloured in salmon. Two views, 90 degrees apart, are shown for clarity.

**Figure S2.**
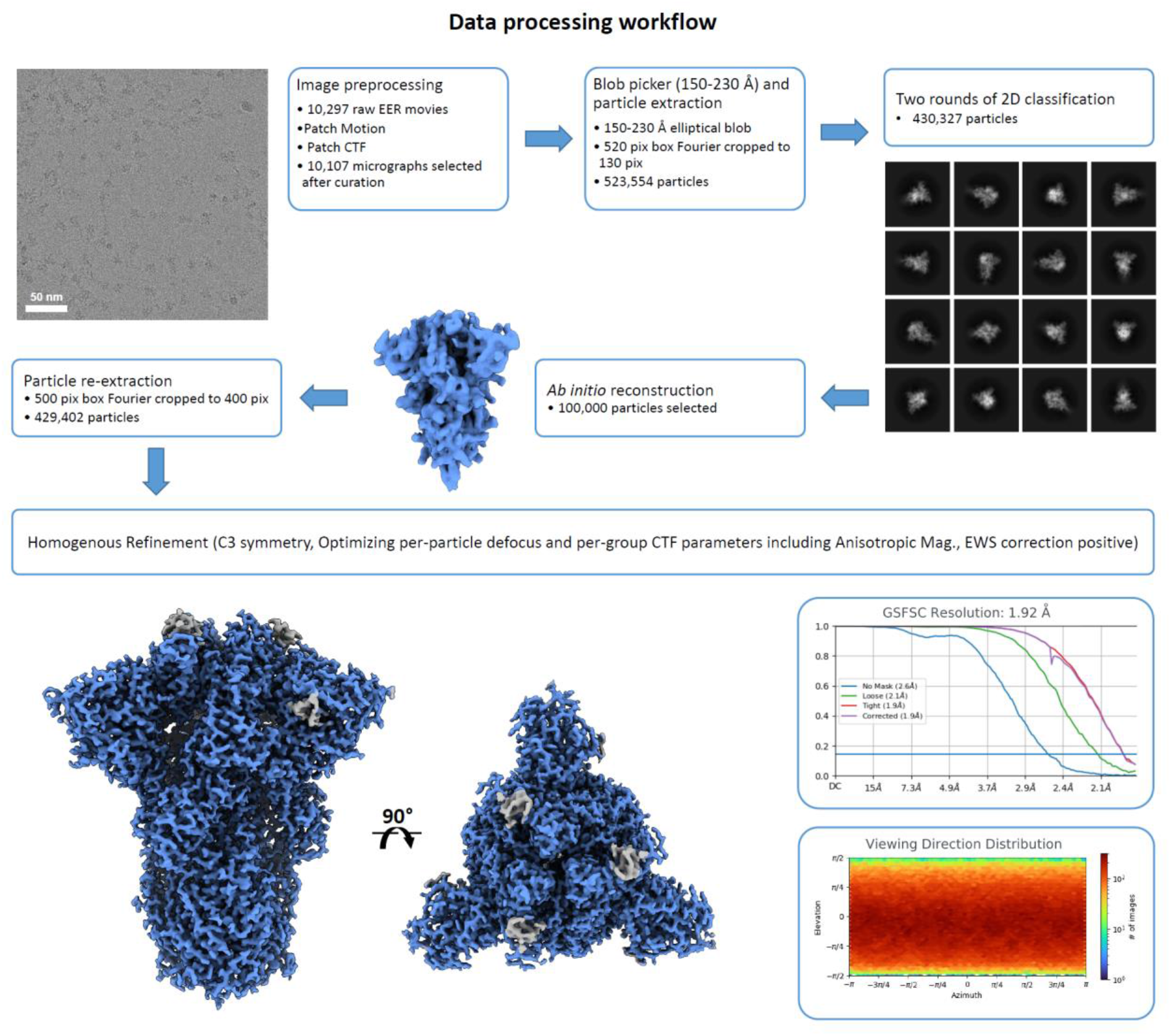
Data processing workflow for Spike S1 in complex with E2E4 biparatopic Bicycle molecule.

**Figure S3.**
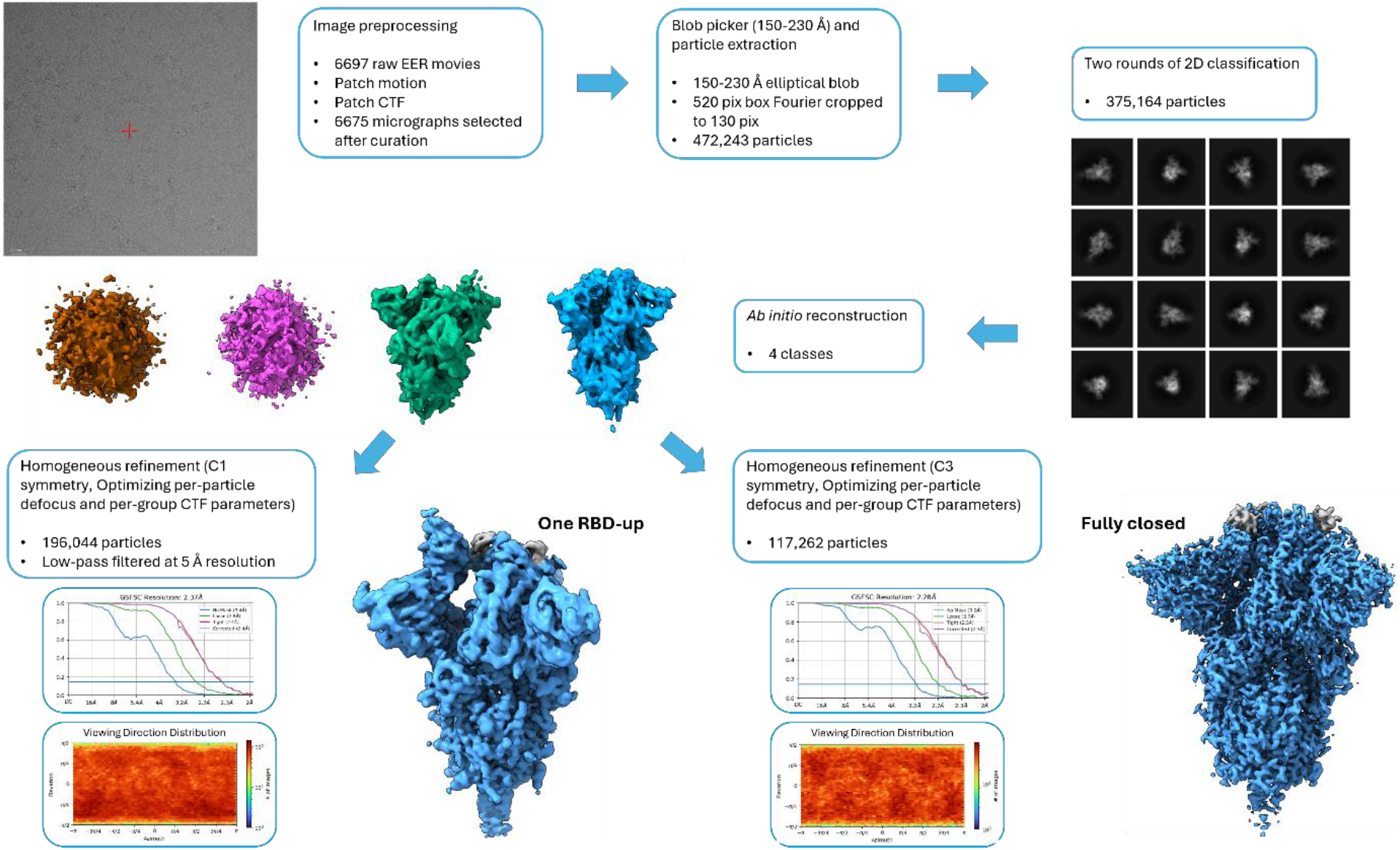
Data processing workflow for Spike S1 in complex with E2 trimer bicycle molecule. The reconstruction of the complex with one “RBD-up” has been low-pass filtered to 5 Å to visualize better the density corresponding to the Bicycle molecules.

**Table S1.** Cryo-EM data collection, model refinement and statistics for Spike S1 in complex with E2E4 biparatopic and E2 trimer.

|  | E2E4 biparatopic<br>(EMD-51660)<br>(PDB: 9GXG) | E2 trimer – one<br>RBD-up<br>(EMD-54164) | E2 trimer -<br>closed<br>(EMD-51659)<br>(PDB: 9GXE) |
| --- | --- | --- | --- |
| **Data collection and processing** |  |  |  |
| Magnification | 165,000 | 165,000 | 165,000 |
| Voltage (kV) | 300 | 300 | 300 |
| Electron exposure (e-/Å<sup>2</sup>) | 50.0 | 59.5 | 59.5 |
| Defocus range (µm) | -0.5 to -1.5 | -0.5 to -1.5 | -0.5 to -1.5 |
| Pixel size (Å) | 0.73 | 0.73 | 0.73 |
| Symmetry imposed | C3 | C1 | C3 |
| Initial particle images (no.) | 523,554 | 472,243 | 472,243 |
| Final particle images (no.) | 429,402 | 196,044 | 117,262 |
| Map resolution (Å) | 1.92 | 2.4 | 2.3 |
| FSC threshold | 0.143 | 0.143 | 0.143 |
| **Refinement** |  |  |  |
| Initial model used (PDB code) | 6XR8 |  | 6XR8 |
| Model resolution (Å) | 2.0 |  | 2.6 |
| FSC threshold | 0.5 |  | 0.5 |
| Map sharpening *B* factor (Å<sup>2</sup>) | 43.9 |  | 45.4 |
| Model composition |  |  |  |
| Non-hydrogen atoms | 28265 |  | 26514 |
| Protein residues | 3371 |  | 3282 |
| Ligands | 97 |  | 57 |
| *B* factors (Å<sup>2</sup>) |  |  |  |
| Protein | 65.5 |  | 98.19 |
| Ligand | 99.6 |  | 117.99 |
| R.m.s. deviations |  |  |  |
| Bond lengths (Å) | 0.006 |  | 0.005 |
| Bond angles (°) | 0.788 |  | 0.739 |
| Validation |  |  |  |
| MolProbity score | 1.34 |  | 1.67 |
| Clashscore | 1.88 |  | 2.81 |
| Poor rotamers (%) | 1.73 |  | 3.13 |
| Ramachandran plot |  |  |  |
| Favored (%) | 96.75 |  | 96.52 |
| Allowed (%) | 3.25 |  | 3.48 |
| Disallowed (%) | 0.00 |  | 0.00 |

**Figure S4.**
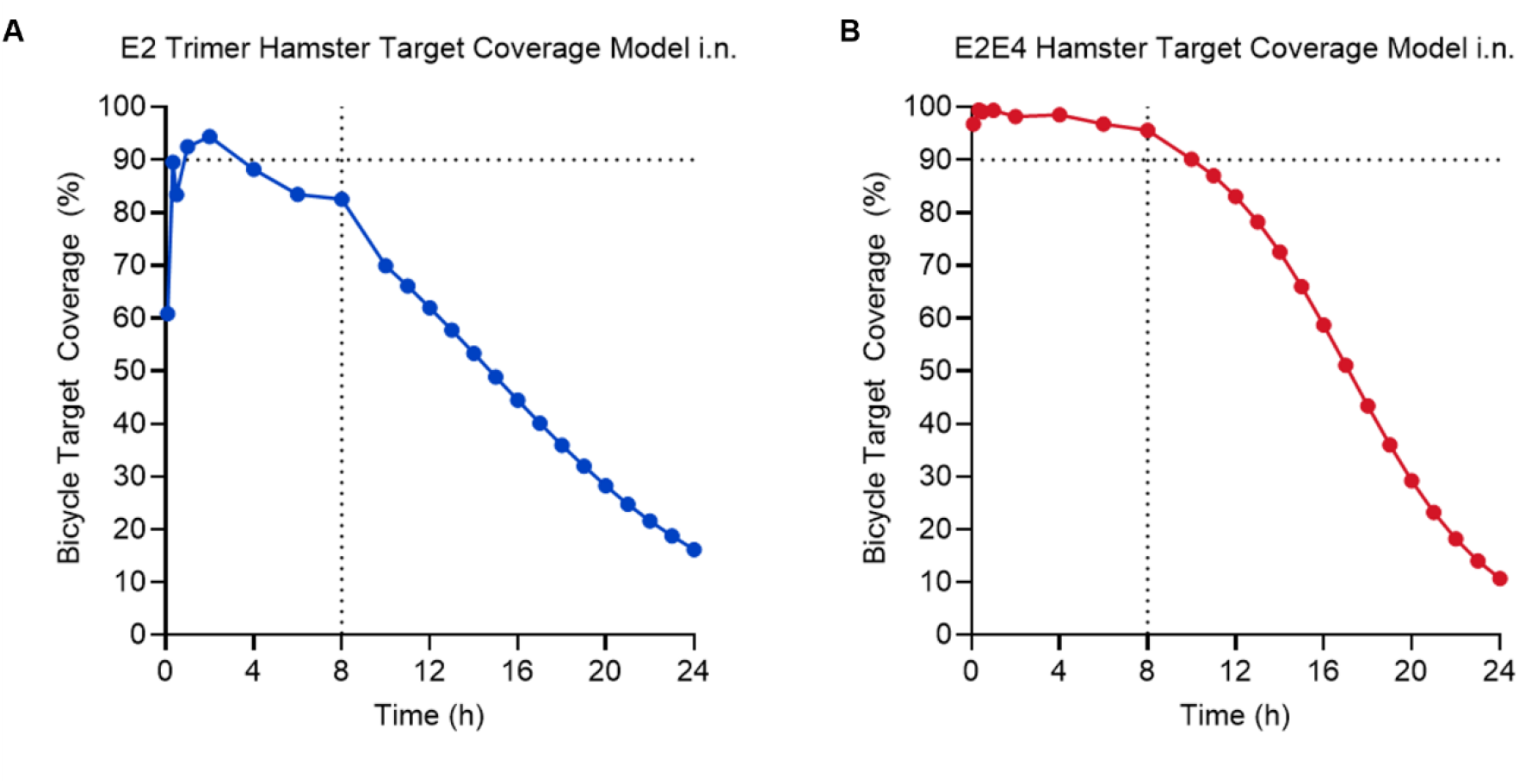
Pharmacokinetic-pharmacodynamic (PKPD) models. PKPD models were used to select an appropriate *in vivo* dosing regimen that would approach 90% target coverage at trough for both the E2 Trimer and E2E4 biparatopic. Using the appropriate data the target coverage predictions for 10 mg/kg are; (A) E2 Trimer hamster i.n. administration (B) E2E4 hamster i.n. administration. Each panel is expressed as % Bicycle Target Coverage over time where black dotted lines reference 90% coverage at 8 hours.

## References

1. Drosten C, Günther S, Preiser W, van der Werf S, Brodt H-R, Becker S, Rabenau H, Panning M, Kolesnikova L, Fouchier RAM, Berger A, Burguière A-M, Cinatl J, Eickmann M, Escriou N, Grywna K, Kramme S, Manuguerra J-C, Müller S, Rickerts V, Stürmer M, Vieth S, Klenk H-D, Osterhaus ADME, Schmitz H, Doerr HW. 2003. Identification of a novel coronavirus in patients with severe acute respiratory syndrome. N Engl J Med 348:1967–76.

2. Zaki AM, van Boheemen S, Bestebroer TM, Osterhaus ADME, Fouchier RAM. 2012. Isolation of a novel coronavirus from a man with pneumonia in Saudi Arabia. N Engl J Med 367:1814–20.

3. Chang C, Ortiz K, Ansari A, Gershwin ME. 2016. The Zika outbreak of the 21st century. J Autoimmun 68:1–13.

4. Mittal N, Medhi B. 2007. The bird flu: a new emerging pandemic threat and its pharmacological intervention. Int J Health Sci (Qassim) 1:277–83.

5. Al Hajjar S, McIntosh K. 2010. The first influenza pandemic of the 21st century. Ann Saudi Med 30:1–10.

6. Mena I, Nelson MI, Quezada-Monroy F, Dutta J, Cortes-Fernández R, Lara-Puente JH, Castro-Peralta F, Cunha LF, Trovão NS, Lozano-Dubernard B, Rambaut A, van Bakel H, García-Sastre A. 2016. Origins of the 2009 H1N1 influenza pandemic in swine in Mexico. Elife 5.

7. Gayle HD, Hill GL. 2001. Global impact of human immunodeficiency virus and AIDS. Clin Microbiol Rev 14:327–35.

8. Zhou P, Yang X-L, Wang X-G, Hu B, Zhang L, Zhang W, Si H-R, Zhu Y, Li B, Huang C-L, Chen H-D, Chen J, Luo Y, Guo H, Jiang R-D, Liu M-Q, Chen Y, Shen X-R, Wang X, Zheng X-S, Zhao K, Chen Q-J, Deng F, Liu L-L, Yan B, Zhan F-X, Wang Y-Y, Xiao G-F, Shi Z-L. 2020. A pneumonia outbreak associated with a new coronavirus of probable bat origin. Nature 579:270–273.

9. Çelik I, Saatçi E, Eyüboğlu AF. 2020. Emerging and reemerging respiratory viral infections up to Covid-19. Turk J Med Sci 50:557–562.

10. Grech AK, Foo CT, Paul E, Aung AK, Yu C. 2024. Epidemiological trends of respiratory tract pathogens detected via mPCR in Australian adult patients before COVID-19. BMC Infect Dis 24:38.

11. Gouglas D, Christodoulou M, Hatchett R. 2023. The 100 Days Mission—2022 Global Pandemic Preparedness Summit. Emerg Infect Dis 29:e221142.

12. Ghattas M, Dwivedi G, Lavertu M, Alameh M-G. 2021. Vaccine Technologies and Platforms for Infectious Diseases: Current Progress, Challenges, and Opportunities. Vaccines (Basel) 9.

13. Feikin DR, Higdon MM, Abu-Raddad LJ, Andrews N, Araos R, Goldberg Y, Groome MJ, Huppert A, O’Brien KL, Smith PG, Wilder-Smith A, Zeger S, Deloria Knoll M, Patel MK. 2022. Duration of effectiveness of vaccines against SARS-CoV-2 infection and COVID-19 disease: results of a systematic review and meta-regression. Lancet 399:924–944.

14. McLean G, Kamil J, Lee B, Moore P, Schulz TF, Muik A, Sahin U, Türeci Ö, Pather S. 2022. The Impact of Evolving SARS-CoV-2 Mutations and Variants on COVID-19 Vaccines. mBio 13:e0297921.

15. Carabelli AM, Peacock TP, Thorne LG, Harvey WT, Hughes J, COVID-19 Genomics UK Consortium, Peacock SJ, Barclay WS, de Silva TI, Towers GJ, Robertson DL. 2023. SARS-CoV-2 variant biology: immune escape, transmission and fitness. Nat Rev Microbiol 21:162–177.

16. Markov P V., Ghafari M, Beer M, Lythgoe K, Simmonds P, Stilianakis NI, Katzourakis A. 2023. The evolution of SARS-CoV-2. Nat Rev Microbiol 21:361–379.

17. Evans RA, Dube S, Lu Y, Yates M, Arnetorp S, Barnes E, Bell S, Carty L, Evans K, Graham S, Justo N, Moss P, Venkatesan S, Yokota R, Ferreira C, McNulty R, Taylor S, Quint JK. 2023. Impact of COVID-19 on immunocompromised populations during the Omicron era: insights from the observational population-based INFORM study. The Lancet Regional Health - Europe 35:100747.

18. Blair HA. 2023. Remdesivir: A Review in COVID-19. Drugs 83:1215–1237.

19. Malin JJ, Weibel S, Gruell H, Kreuzberger N, Stegemann M, Skoetz N. 2023. Efficacy and safety of molnupiravir for the treatment of SARS-CoV-2 infection: a systematic review and meta-analysis. Journal of Antimicrobial Chemotherapy 78:1586–1598.

20. Najjar-Debbiny R, Gronich N, Weber G, Khoury J, Amar M, Stein N, Goldstein LH, Saliba W. 2023. Effectiveness of Paxlovid in Reducing Severe Coronavirus Disease 2019 and Mortality in High-Risk Patients. Clin Infect Dis 76:e342–e349.

21. Williams BA, Jones CH, Welch V, True JM. 2023. Outlook of pandemic preparedness in a post-COVID-19 world. NPJ Vaccines 8:178.

22. von Delft A, Hall MD, Kwong AD, Purcell LA, Saikatendu KS, Schmitz U, Tallarico JA, Lee AA. 2023. Accelerating antiviral drug discovery: lessons from COVID-19. Nat Rev Drug Discov 22:585–603.

23. Gaynor KU, Vaysburd M, Harman MAJ, Albecka A, Jeffrey P, Beswick P, Papa G, Chen L, Mallery D, McGuinness B, Van Rietschoten K, Stanway S, Brear P, Lulla A, Ciazynska K, Chang VT, Sharp J, Neary M, Box H, Herriott J, Kijak E, Tatham L, Bentley EG, Sharma P, Kirby A, Han X, Stewart JP, Owen A, Briggs JAG, Hyvönen M, Skynner MJ, James LC. 2023. Multivalent bicyclic peptides are an effective antiviral modality that can potently inhibit SARS-CoV-2. Nat Commun 14.

24. Shoham S, Batista C, Ben Amor Y, Ergonul O, Hassanain M, Hotez P, Kang G, Kim JH, Lall B, Larson HJ, Naniche D, Sheahan T, Strub-Wourgaft N, Sow SO, Wilder-Smith A, Yadav P, Bottazzi ME, Lancet Commission on COVID-19 Vaccines and Therapeutics Task Force. 2023. Vaccines and therapeutics for immunocompromised patients with COVID-19. EClinicalMedicine 59:101965.

25. Evans RA, Dube S, Lu Y, Yates M, Arnetorp S, Barnes E, Bell S, Carty L, Evans K, Graham S, Justo N, Moss P, Venkatesan S, Yokota R, Ferreira C, McNulty R, Taylor S, Quint JK. 2023. Impact of COVID-19 on immunocompromised populations during the Omicron era: insights from the observational population-based INFORM study. The Lancet regional health Europe 35:100747.

26. Huo J, Mikolajek H, Le Bas A, Clark JJ, Sharma P, Kipar A, Dormon J, Norman C, Weckener M, Clare DK, Harrison PJ, Tree JA, Buttigieg KR, Salguero FJ, Watson R, Knott D, Carnell O, Ngabo D, Elmore MJ, Fotheringham S, Harding A, Moynié L, Ward PN, Dumoux M, Prince T, Hall Y, Hiscox JA, Owen A, James W, Carroll MW, Stewart JP, Naismith JH, Owens RJ. 2021. A potent SARS-CoV-2 neutralising nanobody shows therapeutic efficacy in the Syrian golden hamster model of COVID-19. Nat Commun 12:5469.

27. Cornish K, Huo J, Jones L, Sharma P, Thrush JW, Abdelkarim S, Kipar A, Ramadurai S, Weckener M, Mikolajek H, Liu S, Buckle I, Bentley E, Kirby A, Han X, Laidlaw SM, Hill M, Eyssen L, Norman C, Le Bas A, Clarke J, James W, Stewart JP, Carroll M, Naismith JH, Owens RJ. 2024. Structural and functional characterization of nanobodies that neutralize Omicron variants of SARS-CoV-2. Open Biol 14:230252.

28. Punjani A, Rubinstein JL, Fleet DJ, Brubaker MA. 2017. cryoSPARC: algorithms for rapid unsupervised cryo-EM structure determination. Nat Methods 14:290–296.

29. Meng EC, Goddard TD, Pettersen EF, Couch GS, Pearson ZJ, Morris JH, Ferrin TE. 2023. UCSF ChimeraX: Tools for structure building and analysis. Protein Sci 32:e4792.

30. Emsley P, Lohkamp B, Scott WG, Cowtan K. 2010. Features and development of Coot. Acta Crystallogr D Biol Crystallogr 66:486–501.

31. Liebschner D, Afonine P V, Baker ML, Bunkóczi G, Chen VB, Croll TI, Hintze B, Hung LW, Jain S, McCoy AJ, Moriarty NW, Oeffner RD, Poon BK, Prisant MG, Read RJ, Richardson JS, Richardson DC, Sammito MD, Sobolev O V, Stockwell DH, Terwilliger TC, Urzhumtsev AG, Videau LL, Williams CJ, Adams PD. 2019. Macromolecular structure determination using X-rays, neutrons and electrons: recent developments in Phenix. Acta Crystallogr D Struct Biol 75:861–877.

32. Salzer R, Clark JJ, Vaysburd M, Chang VT, Albecka A, Kiss L, Sharma P, Gonzalez Llamazares A, Kipar A, Hiscox JA, Owen A, Aricescu AR, Stewart JP, James LC, Löwe J. 2021. Single-dose immunisation with a multimerised SARS-CoV-2 receptor binding domain (RBD) induces an enhanced and protective response in mice. FEBS Lett 595:2323–2340.

